# Ferroportin Depletes Iron Needed for Cell Cycle Progression in Head and Neck Squamous Cell Carcinoma

**DOI:** 10.1101/2022.08.26.505485

**Authors:** B. Ross Belvin, Janina P. Lewis

## Abstract

Ferroportin (FPN), the only identified eukaryotic iron efflux channel, plays an important role in iron homeostasis and is down regulated in many cancers. To determine if iron related pathways are important for HNSCC progression and proliferation, we utilize a model of FPN over-expression to simulate iron depletion and probe associated molecular pathways. HNSCC cells are sensitive to iron chelation and ferroptosis, but a non-transformed normal oral keratinocyte (NOK) cell line is not. We found that FPN expression inhibits HNSCC cell proliferation and colony formation but NOK cells are unaffected. Inhibition of cell proliferation is rescued by the addition hepcidin. Decreases in proliferation are due to the disruption of iron homeostasis via loss of labile iron caused by elevated FPN levels. This in turn protects HNSCC cells from ferroptotic cell death. Expression of FPN induces DNA damage, activates p21 and reduces mRNA levels of cyclin proteins thereby inhibiting cell cycle progression of HNSCC cells, arresting cells in S-phase. Induction of FPN severely inhibits Edu incorporation and increases β-galactosidase activity, indicating cells have entered senescence. Finally, in an oral orthotopic mouse xenograft model, FPN induction yields a decrease of tumor growth. Our results indicate that iron plays a role in HNSCC cell proliferation and sustained growth and ferroptosis iron based therapeutic strategies may have potential therapeutic benefit.

## Introduction

Head and neck squamous cell carcinoma (HNSCC) is the 6^th^ most common malignancy worldwide with more than 800,000 new cases and 450,000 deaths annually (1). Patients with recurrent or metastatic (R/M) HNSCC have a poor prognosis, with a median survival rate of under 1 year (2). Due to this, there is an ongoing effort to better understand the drivers of HNSCC progression and metastatic spread to develop novel and effective therapeutic strategies (3,4). Iron has emerged a key metabolic driver of malignancy, with many cancer types exhibiting an iron “addicted” phenotype.

The increased acquisition of iron in tumor cells has been shown to contribute to their growth and progression (5). Due to their increased proliferation, cancer cells have an elevated metabolic demand for iron when compared to normal cells (6). Consequently, genes related to iron homeostasis are commonly deregulated in many cancer types. Through alterations in cellular iron metabolism, individual cancer cells accumulate higher levels of labile iron through mechanisms involving increased iron uptake and retention as well as decreased iron efflux (7). However, our understanding of iron homeostasis in HNSCC is scarce.

It is noteworthy that eukaryotic cells utilize multiple mechanisms to import iron, but there is only one mechanism to export iron out of cells. This mechanism utilizes the iron efflux channel ferroportin (FPN) (8). Although it is of significant importance in many malignancies and iron-based disorders, the role of FPN in oral keratinocytes is yet to be determined. The repression of FPN expression has been demonstrated in multiple cancer types and provides a mechanism for retention of cellular bio-available iron needed for cancer cell proliferation and cell cycle progression (9–11). However, this increased intra-cellular iron also makes cancer cells vulnerable to iron-based therapies such as ferroptosis and iron chelation (12,13).

There is growing evidence that iron metabolism is altered in HNSCC. Mechanisms involving both an increase in iron uptake and retention have been observed. There is a noted increase in the levels of transferrin receptor 1 (TFR1) in clinical isolates of esophageal tumors (14). Furthermore, co-amplification of *TFRC* and *PIK3Ca* genes correlated with increased distant metastasis and poor prognosis in HNSCC patients (15). There is mounting evidence that the iron storage protein, ferritin (FTN), is significantly up-regulated in HNSCC and this increase corresponds to poorer prognosis (16–18). Furthermore, increased levels of serum ferritin correlate with an increase in HNSCC lymph node metastasis (19). In a panel of HNSCC cell lines, homeostatic iron regulator protein (HFE) over-expression increased intra-cellular iron via an increase in hepcidin expression (20). Hepcidin binds directly to FPN, thereby causing its internalization and eventual degradation (21).

However, despite the evidence of deregulation of iron related genes in HNSCC little is known about the molecular mechanisms behind the abnormality. Specifically, there are no studies focused on the impact FPN has on HNSCC progression. Even less is understood why HNSCC cells deregulate their iron homeostasis. Studies in other cancer types have implicated elevated iron levels accounting for increased activity of iron containing enzymes involved in DNA replication and nucleotide synthesis (22,23). In this study, we modulate FPN levels to better understand the function of iron in advanced, metastatic HNSCC cell lines to understand how loss of iron affects growth and proliferation in these cells. We reveal that FPN mediated iron efflux has substantial effects on HNSCC growth and is capable of inducing DNA damage, cell cycle arrest, and subsequent senescence in these cells but not in a non-malignant, tert-immortalized normal oral keratinocyte (NOK) model. Ultimately, expression of FPN negatively effects tumor growth and progression in a murine model of oral cancer.

## Results

### HNSCC cells exhibit increased iron dependence and are susceptible to ferroptosis

Iron based therapies are emerging as a potential therapeutic strategy for the treatment of HNSCC. We tested the susceptibility of 4 HNSCC cell lines derived from metastatic sites to iron chelators DFO and Dp44mT (24,25). As a non-malignant control, a TERT-immortalized non-transformed normal oral keratinocyte (NOK) cell line was used (Fig. 1 A-B). The HNSCC cells showed significantly higher sensitivity to iron chelation when compared to the non-transformed NOK cell line.

**Figure 1.**
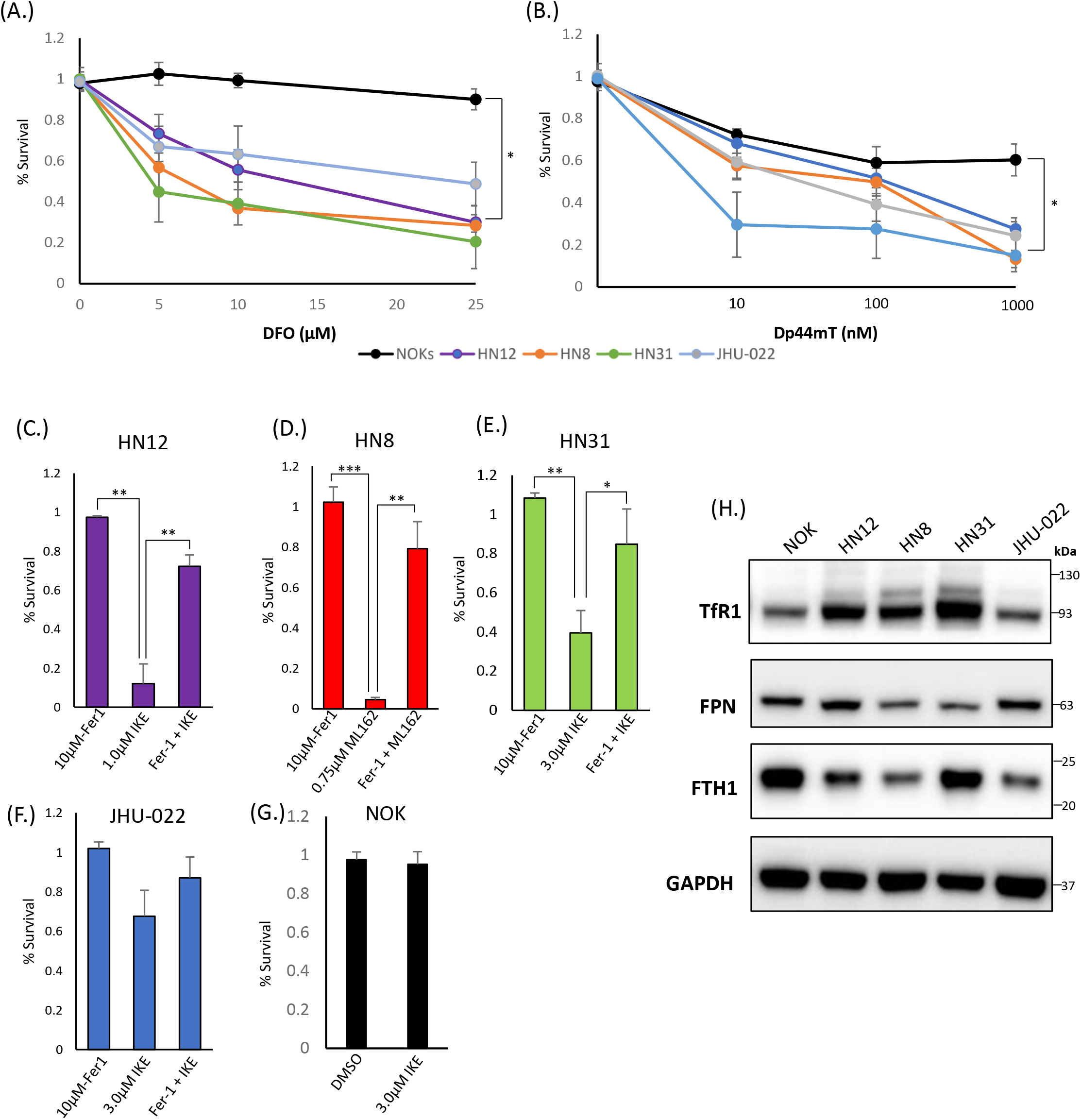
HNSCC cell lines exhibit increased dependence on iron and are susceptible to iron chelation and ferroptosis. **A**. Cells were grown with increasing amounts of deferoxamine (DFO) for 72 hours and cell viability was assessed via CellTiter-glo assay. **B**. Cells were treated with the indicated amount of Dp44mT for 72 hours and cell viability was measured via CellTiter-glo. **C-G**. The indicated cell line was seeded and treated with Ferrostatin-1 (10μM), the indicated ferroptosis inducer (IKE or ML162), or a combination of both for 72 hours. Cell viability was measured via CellTiter-glo assay. Values were normalized to a non-treated control with DMSO. **H**. Western Blot of TFR1, FPN, FTH1, with GAPDH as a loading control. (*, P<0.05; **,P<0.01; ***,P<0.001)

Next we tested the susceptibility of these cell lines to ferroptosis inducers Imidazole Ketone Erastin (IKE) and ML162 (Fig. 1 C-G; Fig. S1) IKE is an analog of erastin that inhibits system xC; ML162 is a potent inhibitor of GPX4. The HN12, HN8, and HN31 cell lines showed significant susceptibility to ferroptosis induced by either IKE or ML162 and were efficiently rescued from cell death by the supplementation with Ferrostatin-1. The JHU-022 cell line showed moderate susceptibility to IKE at the highest concentration (3 μM). As with the iron chelators, the NOKs showed little to no decrease in cell viability in response to IKE or ML162.

To corroborate these data, we next looked at TFR1, FPN, and ferritin heavy chain (FTH1) protein levels in these cell lines to get a snapshot of iron homeostasis. We found that, consistent with previous published data, the TFR1 levels were higher in the HN12, HN8, and HN31 cell lines when compared to a non-malignant control. This corresponded to the cells most susceptible to ferroptosis inducers. FPN was noticeably lower in the HN8 and HN31 lines as well. The JHU-022 line, which was most resistant to ferroptosis of the cancer cell lines, had the lowest levels of TFR1. The non-malignant NOK cell line had much lower TFR1 and high FPN levels. NOK cells had the highest levels of FTH1 and the HNSCC lines had much lower FTH1 levels.

### FPN over-expression disrupts iron metabolism HNSCC

To determine if changes in iron metabolism are a potential driver of HNSCC growth and proliferation or merely incidental we utilized an FPN over expression system and assessed its effects on HNSCC cell lines. The FPN CDS was cloned into the pLVX-TetOne-puro vector to create a conditional doxycycline-driven system to express FPN. The pLVX-Tet FPN and control vectors (pLVX-Tet Luc) were stably transduced into the HN12, JHU-022, and NOK cell lines (Fig. 2A). As a control, cells treated with doxycycline were concurrently treated with hepcidin. Hepcidin successfully knocked down the protein levels of FPN indicating that FPN was at the cell surface and active.

**Figure 2.**
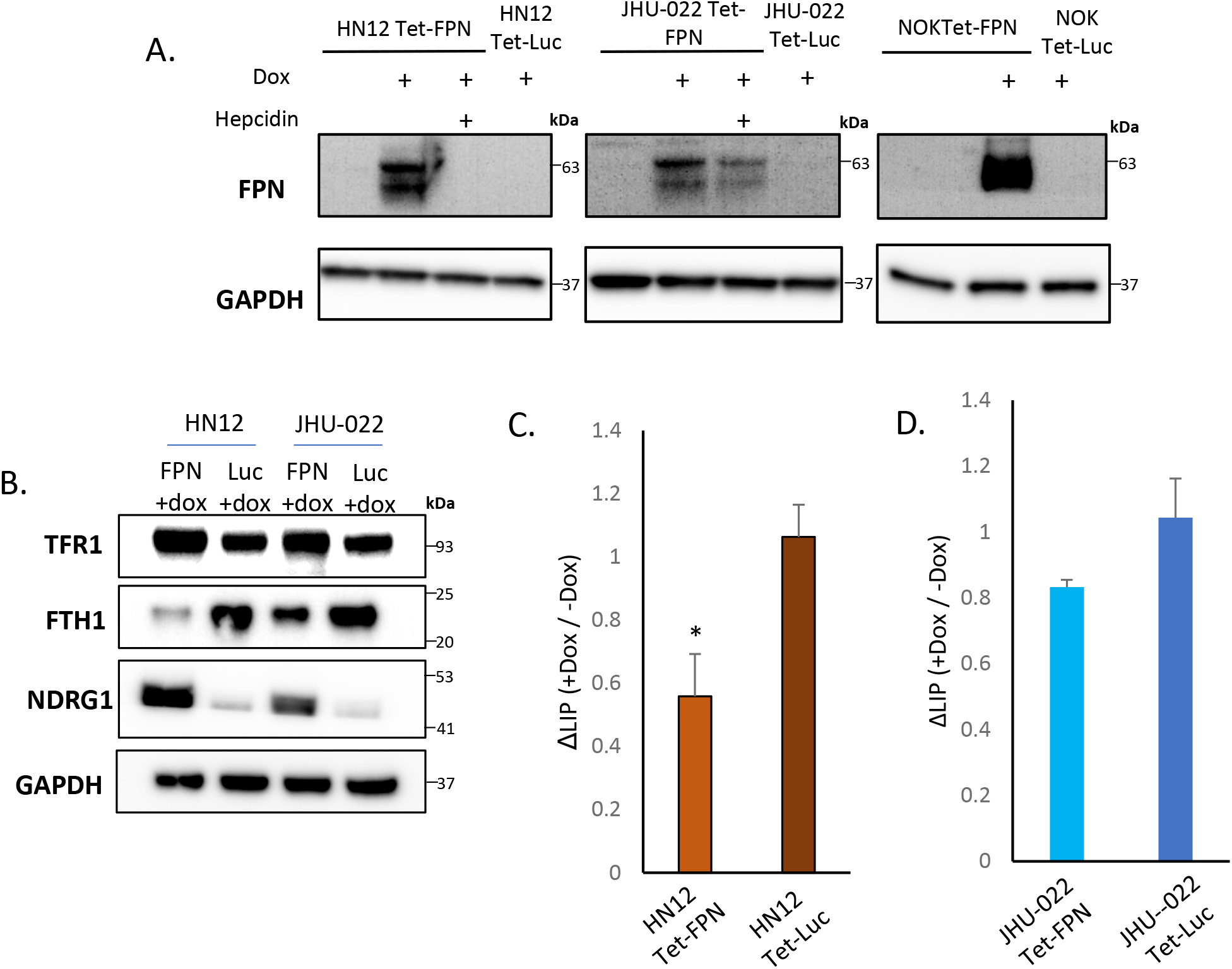
Stable Expression of FPN in the HN12, JHU-022, and NOK cell lines. **A**. Western blot of FPN in the HN12-FPN/Luc, JHU-022-FPN/Luc, and NOK-FPN/Luc after treatment with 0.5 μg/mL of doxycycline for 48 hours. For rescue experiments, hepcidin was added to cell cultures with doxycycline at 20μM. **B**. HN12-FPN/Luc and JHU-022-FPN/Luc were stimulated with 0.5 μg/mL dox for 72 hours and blotted for iron related proteins transferrin receptor 1 (TfR1), ferritin heavy chain 1 (FTH1), or N-myc downstream regulated protein (NDRG1). **C,D**. Measurement of the labile iron pool in cells (*, P<0.05). All data is representative of 3 biological replicates.

FPN drives cellular iron efflux, thus FPN expression should lead to a decrease in the labile iron levels. This change should be reflected in the regulation of iron related genes. Consistent with this prediction, the HN12-FPN and JHU022-FPN lines had elevated levels of TfR1 and decreased levels of ferritin heavy chain 1 (FTH1) when compared to Luciferase expressing controls (Fig 2B). NDRG1 (a gene commonly viewed as a metastasis suppressor) has been showed to be elevated in iron deplete conditions (26). We found the levels of NDRG1 to be highly elevated in the FPN expressing lines (Fig. 2B). To substantiate this, we directly measured the levels of labile iron using the FerroOrange dye. We found that levels of labile iron were decreased in the FPN expressing lines when compared to controls (Fig. 2C-D).

### FPN expression inhibits HNSCC cell growth and proliferation

We tested whether FPN mediated iron depletion effected cell proliferation. We found that expression of FPN caused significant decrease in growth in the HNSCC cell lines (HN12 and JHU-022) when compared to a luciferase expressing control cell line when assessed via cell counts, Celltiter blue assay and clonogenic assays (Fig. 3 A-F). After 4 days, growth of the FPN expressing HNSCC cell lines were significantly lower than their Luciferase expressing control lines. In the HN12 line specifically, the significant differences in growth were seen after 48 hours, with little to no growth occurring after this time point (Fig. 3A). The non-transformed NOK cell line showed no significant difference in growth between the FPN expressing line and luciferase control over 4 days (Fig 3G-H). These results were also confirmed utilizing a clonogenic assay (Fig. 3I)

**Figure 3.**
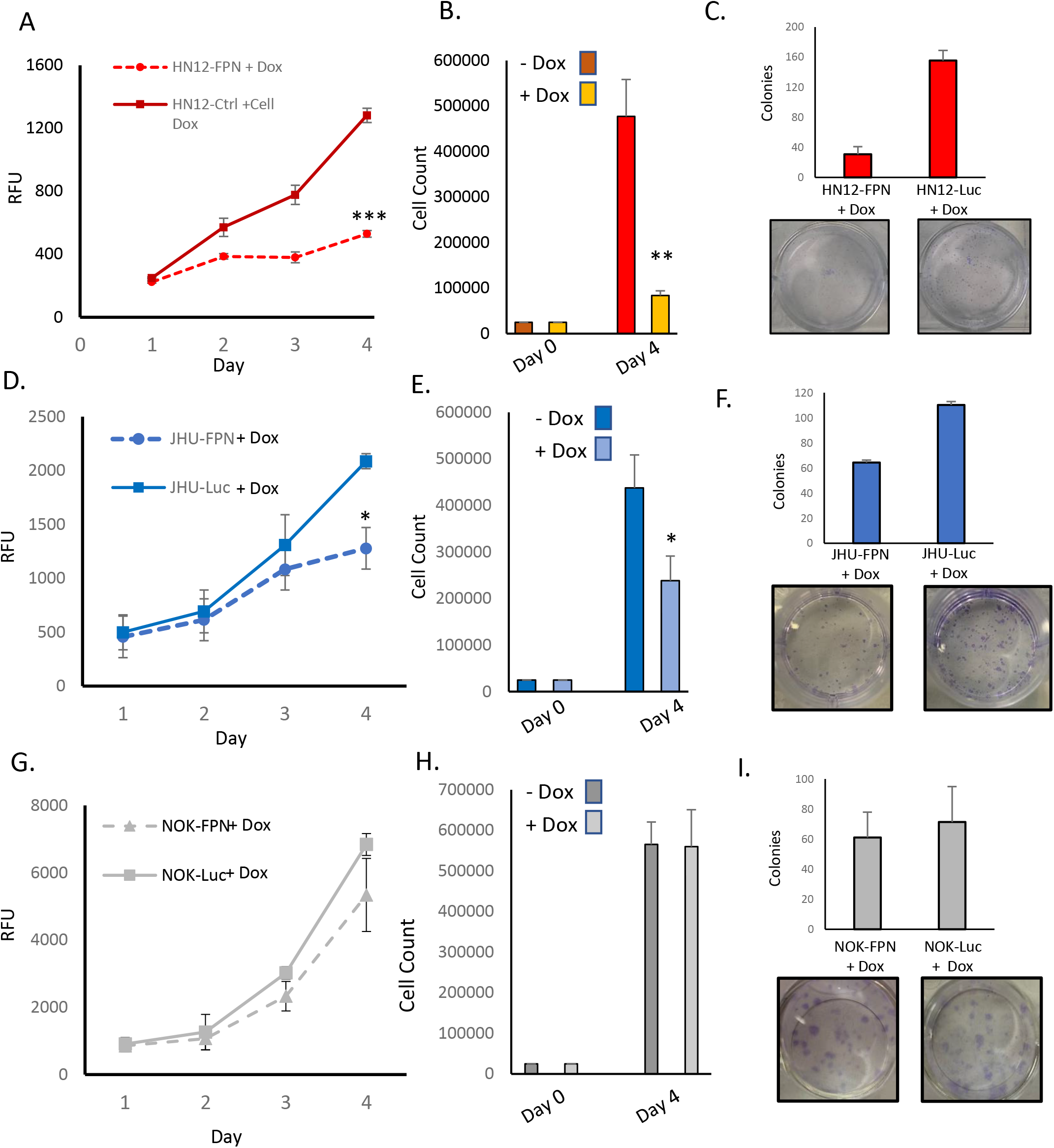
Expression of FPN inhibits growth and proliferation of HNSCC. **A, D, G**. HN12 (A), JHU-022 (D) and NOK (G) FPN expressing or Luc control expressing cells were seeded in 96 well plates at 1000 cells per well with 0.25 μg/mL dox and grown for 4 days. Growth was assessed using CellTiter-blue reagent. **B, E, H**. HN12-FPN (B), JHU-022-FPN (D), and NOK-FPN (H) cells were seeded at 25,000 cells per well in a 6 well plate +/−0.25 μg/mL dox. After 4 days cells were counted via hemocytometer. **C, F, I**. Clonogenic assay quantification and representative plate of HN12 (C), JHU-022 (F), and NOK (I) FPN expressing or Luc control expressing cells. Cells were seeded at 250 cells per well with 0.25 μg/mL dox and colonies were counted after 7-8 days. (*, P<0.05; **,P<0.01; ***,P<0.001). All data is representative of 3 biological replicates.

To ensure that FPN was expressed on the cell membrane and that the phenotype observed was due to active FPN we treated the HNSCC cell lines with doxycycline and hepcidin. We found that hepcidin rescued the growth of the HNSCC cell lines when assessed via cell counts (Fig. S2).

We have shown that the HN12 cell line is sensitive to ferroptosis induced by IKE treatment. Ferroptosis is dependent on elevated iron levels to induce lipid peroxide damage. Decreasing iron levels protect cells from ferroptotic cell death. Consistent with this, expression of ferroportin efficiently rescues the HN12 cell line from ferroptosis induced cell death (Fig. S3).

### FPN expression inhibits HNSCC cell cycle progression

Iron levels are linked to progression of the cell cycle (27,28). To determine the mechanisms by which FPN expression inhibits HNSCC growth and proliferation, we assessed the cell cycle of FPN expressing cell lines. As seen in Figure 4, FPN expression in HNSCC cell lines triggered cell cycle arrest in the S-phase. Specifically, most HN12-FPN cells were arrested in the S-phase (Fig 4A). The FPN expression in the JHU022 line yielded a modest arrest in the S-Phase (Fig. 4B). FPN expression had no significant effect on the cell cycle in the NOK cell line (Fig. 4C). Interestingly, this is not the case with cells treated with the iron chelator DFO (Fig S4). When treated with 50 uM DFO, HN12 cells are arrested in the G1 phase. This is consistent with previous studies: Iron chelation typically causes cell cycle arrest in the G1 phase. Thus, a different mechanism may be at play in the FPN induced cell lines.

**Figure 4.**
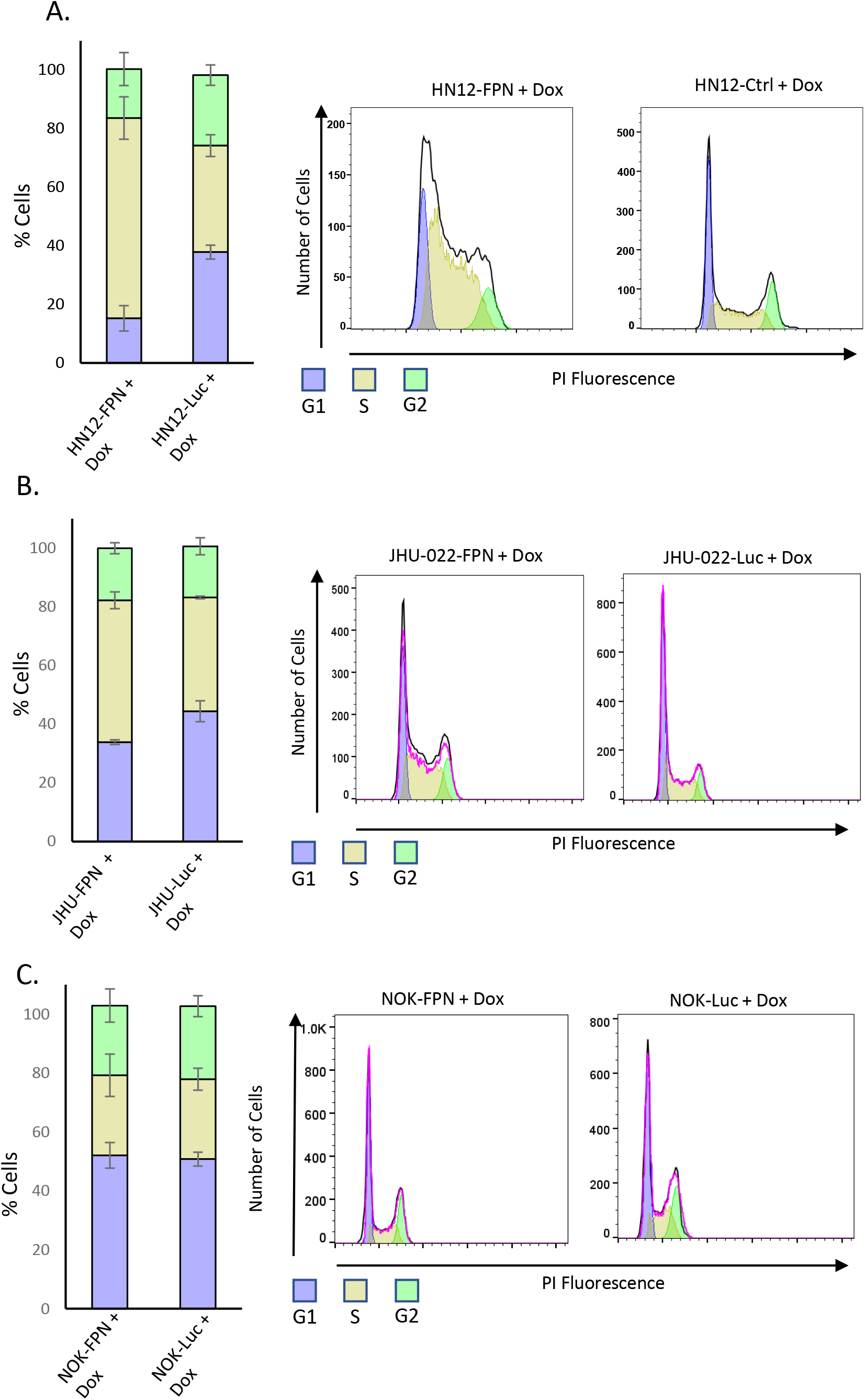
FPN expression causes cell cycle arrest in HNSCC. FPN or Luciferase control expressing HN12 (**A**.), JHU-022 (**B**.) and NOK (**C**.). were grown with 0.5 μg/mL dox for 3 (HN12) or 4 days (NOK, JHU-022) and stained with propidium iodide to measure DNA content. The DNA content was assessed by flow cytometry and the cell cycle distribution was analyzed using FlowJo. All data is representative of 3 biological replicates.

We next evaluated the effect FPN expression on key cell cycle factors. The induction of p21 is a major S phase checkpoint pathway, thus we suspected the p21 may be responsible for the S phase arrest seen in the HNSCC cell lines. We found that the levels of p21 were significantly elevated in the HN12-FPN line at the RNA and protein level (Fig. 5 A-B) but not in the JHU-022-FPN line. (Fig. 5C). Consistent with this, we found that the transcript levels of cyclins A, B, and D were all significantly decreased in the HN12-FPN line. In the JHU-022-FPN line, we found only Cyclin A and D were down regulated (Fig 5C).

**Figure 5.**
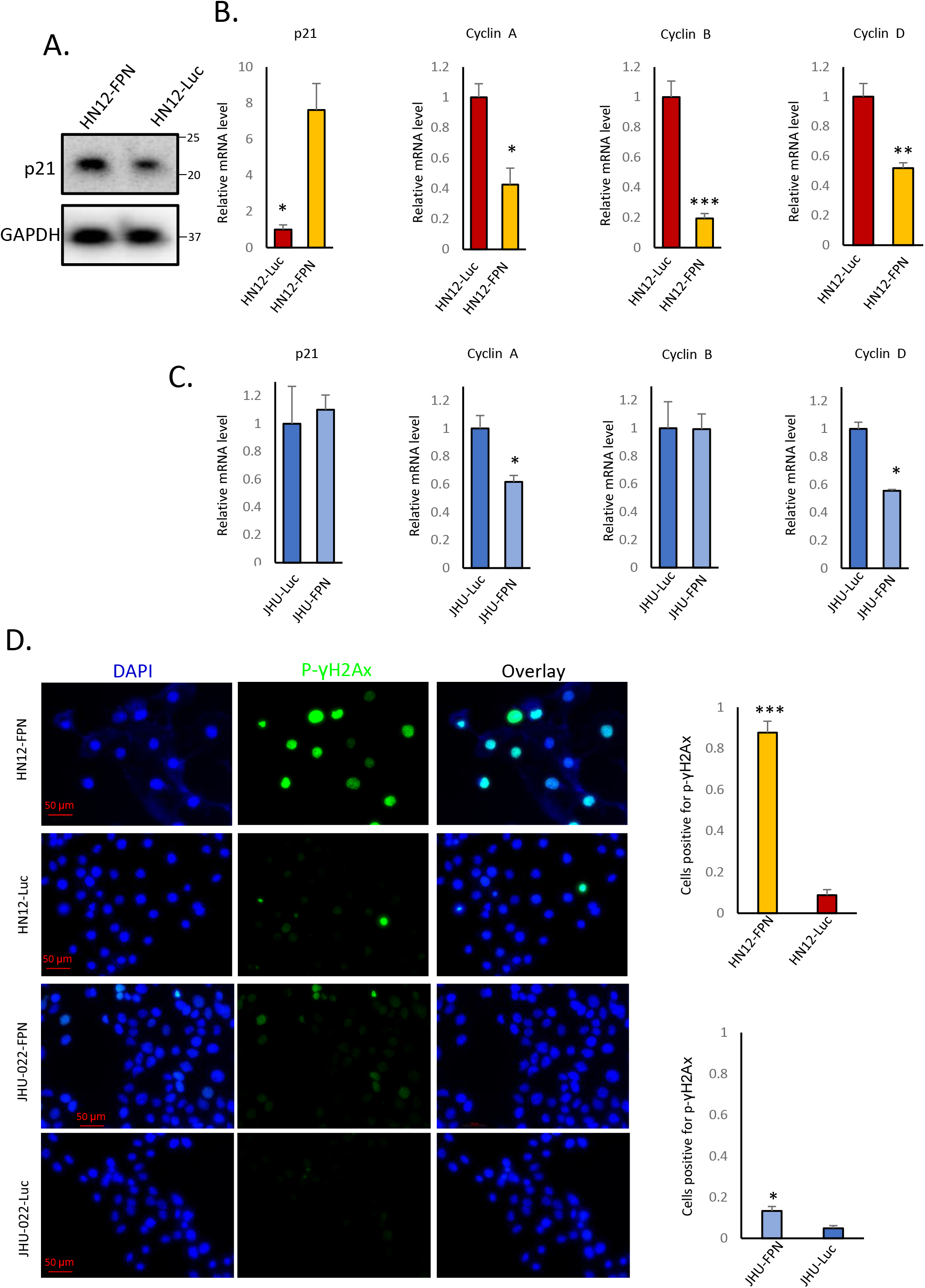
FPN expression in HNSCC cell lines downregulates key cell cycle related genes. **A**. Western blot of p21 levels in HN12-FPN and HN12-Luc expressing cells. **B, C**. The mRNA levels of p21, cyclin A, cyclin B, and cyclin D were assessed by qRT-PCR in HN12-FPN/Luc and JHU-022-FPN/Luc cells grown in 0.5 μg/mL dox for 3 (HN12) or 4 (JHU-022) days. Levels of target genes are normalized to the levels of both GAPDH and beta-actin. Data is representative of 3 techincal replicates performed in 3 biological replicates. **D**. HN12-FPN/Luc and JHU-022-FPN/Luc cells stimulated with 0.5 μg/mL dox for 3 days and stained for p-γH2Ax (green) or DAPI (blue). Images are representative of 3 biological replicates. (*, P<0.05; **,P<0.01; ***,P<0.001)

Since p21 is induced by DNA damage and cyclins are down regulated by DNA damage, we asked whether FPN expression and iron depletion led to the accumulation of DNA damage. Phosphorylation of histone gamma H2AX is recognized marker for DNA double strand breakers. As seen in Fig. 5D, FPN overexpression led to a large increase in the levels of phospho(S139)-γH2AX in HN12-FPN cells. Comparatively, there was a modest increase in phospho(S139)-γH2AX in the JHU-022-FPN cells.

### FPN expression induces senescence in the HN12 cell line

Senescence is a well-documented response in cells experiencing catastrophic DNA damage. To determine whether FPN expression induced senescence in HNSCC we utilized an Edu incorporation assay to determine if DNA synthesis and replication is attenuated. Doxycycline induced FPN expression significantly reduced Edu incorporation in the HN12 cell line (Fig. 6A). Loss of Edu incorporation is consistent with cells under a senescent phenotype. Despite seeing a modest increase in the S phase of the cell cycle in the JHU-022 cell line, FPN expression does not yield a significant change in the Edu incorporation during the experimental time frame (1.5 hours) (Fig. 6B).

**Figure 6.**
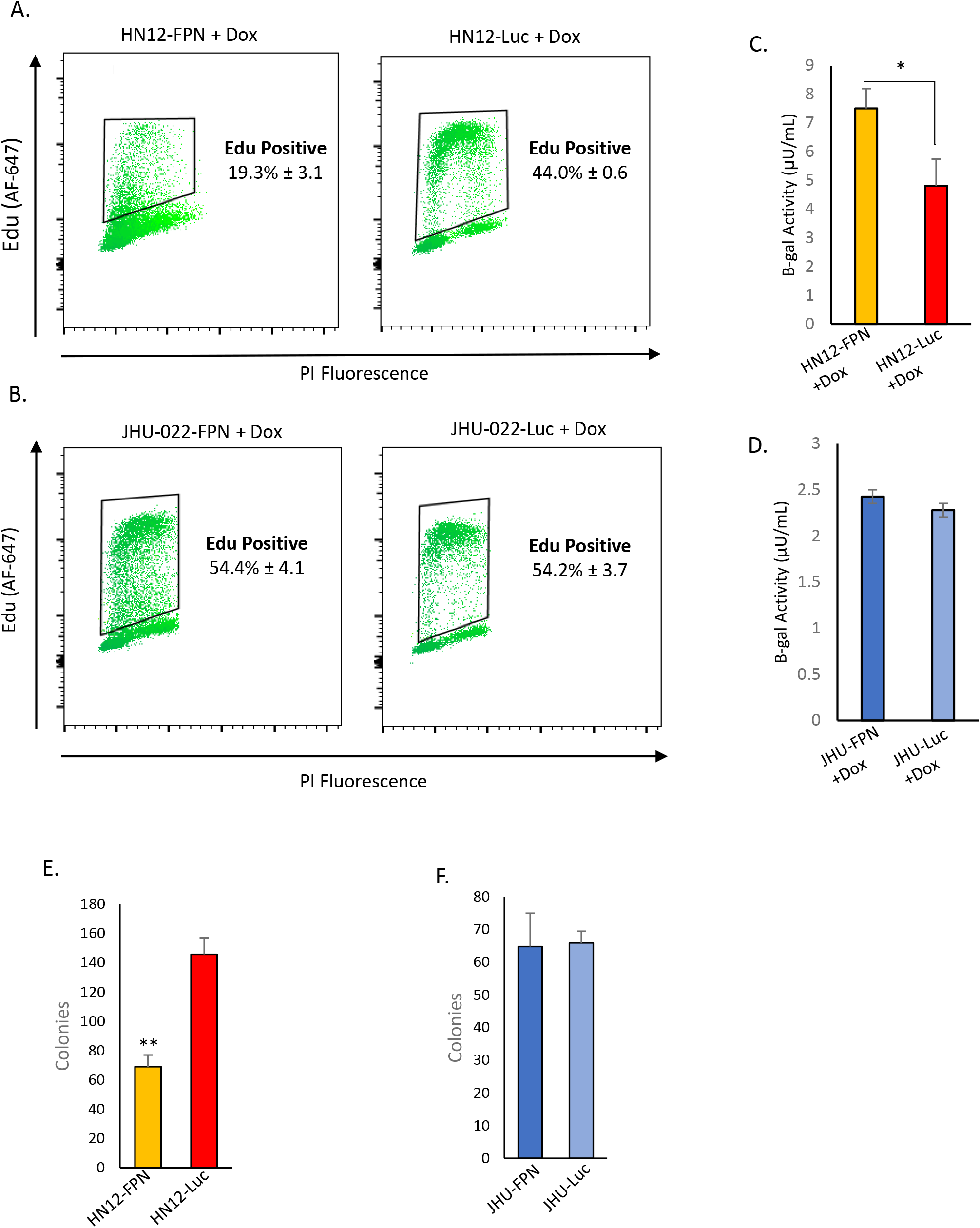
FPN expression causes HN12 cells to become senescent. **A, B**. EDU incorporation of HN12-FPN/Luc (A) and JHU-022-FPN/Luc (B) grown with 0.5 μg/mL dox for 3 days. **C,D**. Beta-galactosidase activity of HN12-FPN/Luc (C) and JHU-022-FPN/Luc (D) after 7 days of exposure to 0.5 μg/mL dox. One μUnit is equal to 1 pmol of product generated per min. **E, F**. Clonogenic assay of HN12 and JHU-022 FPN and Luciferase expressing grown for 72 hours with 0.25 μg/mL dox for 72 hours, then recovered with dox free media for 48 hours.

Senescence associated beta-galactosidase activity is one the most widely used biomarkers to determine the state of a cell (29). To confirm that FPN expression is inducing senescence in the HN12 cell line and not the JHU-022 cell line, we measured the B-galactosidase activity. We found that the B-gal activity is significantly higher in the HN12-FPN induced cell line when compared to the Luc expressing control (Fig. 5C). Consistent with previous data, there is no difference in the B-gal activity in the JHU-022 cell line (Fig. 5D). One of the most recognizable properties of senescence is the enlarged and flattened morphology of these cells. HN12 cells adopt a senescent cell morphology when FPN expression is induced (Fig. S5).

To determine if FPN induced senescence was transient, HN12 and JHU-022 FPN and luciferase expressing lines were grown with doxycycline for 3 days. The doxycycline was removed and cells were washed and grown for additional 2 days with fresh dox free media to recover. A clonogenic assay was performed on these cells. We found that the HN12-FPN cell line still had significantly lower number of colonies recovered (Fig. 6E) indicating the FPN mediated senescence was persistent. There was no difference in the colony numbers of the JHU-022-FPN and JHU-022-Luc lines after recovery in dox free media (Fig. 6F).

Finally, we wanted to test if deoxynucleotide depletion contributed to the senescent phenotype seen in the HN12-FPN cells. Studies have revealed that iron plays a crucial role in the maintenance of the dNTP pool needed for DNA synthesis as a component of ribonucleotide reductase (30–32). We supplemented HN12-FPN cells with deoxynucleotides and assessed their growth after 48 hours (Fig S6). We saw a partial rescue of the growth of these cells, however, over longer time courses this effect was not sustained.

### FPN expression inhibits growth of HNSCC tumors in an orthotopic xenograft model

Given that FPN expression can induce cell cycle arrest and inhibit cancer cell proliferation, we utilized an oral orthotopic xenograft model to test growth of the HN12 cell line *in vivo*. HN12-FPN cell line was implanted into in the left cheek of NSG mice. After tumors had grown for 5 days, mice were randomized into two groups. Doxycycline was added to the drinking water to induce FPN expression in the experimental group and the control group was provided drinking water without doxycycline. In total, tumors were grown for 21 days before termination of the experiment due to morbidity of mice in the control group. Tumors were harvested, and tumor volume and weight were assessed. Measurements indicate tumor weight and volume were significantly lower in the experimental group (FPN induced) when compared to the control group (Fig. 7A-B). These results indicate the FPN expression suppresses growth of HNSCC *in vivo*.

**Figure 7.**
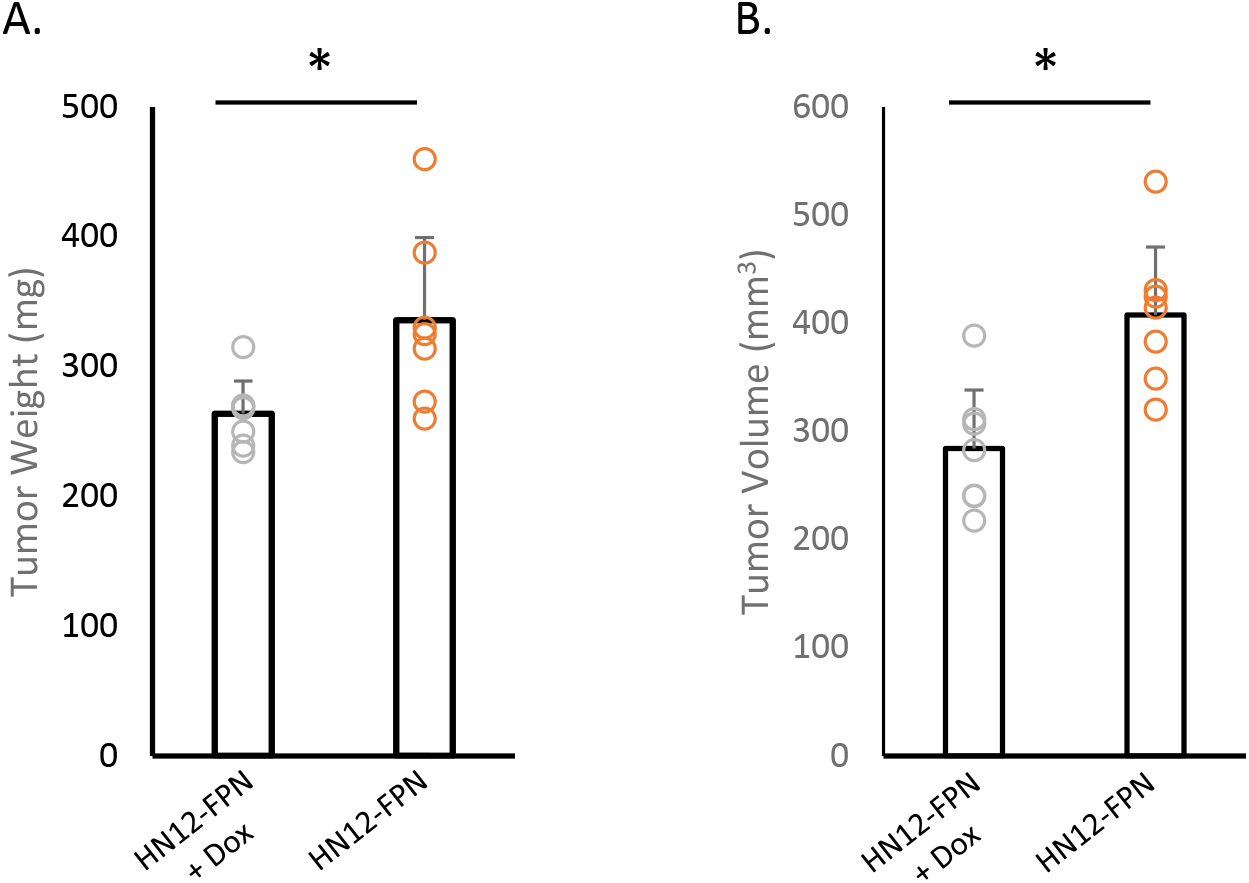
FPN expression inhibits growth of HN12 cells in an orthotopic xenograft model. HN12-FPN cells were implanted into the left cheek of mice and allowed to form tumors for 5 days. FPN expression was induced by adding 2mg/mL Doxycycline to the drinking water of mice. Control mice received normal drinking water. After 21 days, tumors were harvested and tumor weight (**A**.) and tumor volume (**B**.) were measured. For each group n=7 mice.

## Discussion

This is the first study to investigate the effect of FPN expression on oral keratinocytes and HNSCC. We show that FPN mediated iron depletion has significant effect on growth inhibition, cell cycle arrest, and senescence. We saw two distinct responses in the HN12 and JHU-022 cell lines. The HN12 cell line exhibited a very iron “addicted” phenotype, where loss of available iron led to near total growth arrest, DNA damage, and senescence. This decrease in proliferation carried over into our *in vitro* orthotopic mouse model: tumors were significantly smaller in the FPN expressing xenografts. The JHU-022 line saw a significant decrease in growth, however, this did not lead to high levels of DNA damage and the cells did not enter senescence. Notably in the HN12 cell line, growth arrest continued despite the removal of doxycycline as assessed by clonogenic assays. Thus, there are still fundamental questions regarding the pathways active during iron depletion in HNSCC.

The addition of deoxynucleotides had partial rescue effect on the HN12-FPN expressing cell line. Previous studies have shown that sufficient iron levels are extremely important for maintain the available pool of deoxynucleotides needed for DNA synthesis and replication (32,33). Lack of available dNTPs for DNA synthesis can lead to cell cycle arrest and DNA damage seen in HN12-FPN cells. A partial rescue was seen over 48 hours, however the addition deoxynucleotides alone was not able to fully rescue the cells from growth inhibition, implying that elevated iron levels may be playing a larger role than maintenance of the dNTP pool in these cells.

FPN expression had little to no effect on the non-malignant NOK cells. This was not unexpected, as the NOKs were resistant to growth inhibition by DFO and cell death via ferroptosis at the concentrations used in this study. Expression of FPN caused significant changes in metabolism in the HNSCC cell lines. Iron homeostasis is regulated by a sensitive feedback system. When cells are depleted of iron, we saw levels of TFR1 go up and levels of FTH1 go down to compensate for the lack of available iron, validating our experimental model.

It should be noted that we made many attempts at creating a stable expression of FPN in the HN8 and HN31 cells, yet we were unsuccessful. When we were able to see FPN expression, the elevated levels were quickly lost after a passage, even after isolation of a monoclonal culture. Notably, these two cell lines have the lowest levels of FPN of the cells we tested. There appears to be a significant selective pressure against FPN expression in these cell lines.

We chose to use a model of FPN expression to simulate iron depletion rather than the normal iron chelation strategies in wide use. FPN is unable to liberate bound iron from non-physiological targets. DFO, and other iron chelators, are capable of stripping iron from bound proteins and storage molecules due to their extremely high affinity for iron (13,34). Iron chelators are well known for their use in exploring the biological mechanisms of iron homeostasis and their therapeutic benefits in iron related pathologies. However, there are noted downsides to iron chelation therapy, which are prone to off target side effects (35). Furthermore, their ability to chelate intra-cellular iron has brought into question if they truly mimic physiologically relevant iron deplete conditions (36). Thus our strategy of FPN expression more resembles an iron starvation phenotype in an iron deplete environment.

Still, there are similarities in the response of these cells to iron chelator treatment and FPN over-expression. The inhibition of cell proliferation, induction of DNA damage, and cell cycle arrest have been observed in DFO treatment. There are also differences. Treatment of cells with DFO causes cells to arrest in G1 phase, via the action of IRP2 (37). In our study, we saw a decrease in the population of cells in G_0_/G_1_ indicating the mechanisms between iron chelation treatment and FPN mediated iron depletion may be different.

Iron is a necessary nutrient for cellular health however, not every cell has the same need for iron. In this study we reveal 3 very different responses to FPN mediated iron depletion. HN12 cells are almost completely incapable of growing without elevated iron levels and enter a senescent state. Even after removal of doxycycline and recovery, their growth is still arrested (Fig. 6E). JHU-022 cells grow at a decreased rate, however, they do not enter senescence and growth resumes immediately after removal of FPN inducer (Fig. 6F). NOK cells are unperturbed by increased FPN expression – they may be able to compensate for this lack of iron in other ways. This implies that different malignant cell lines have different requirements for iron supplementation. What drives this difference, and how this may be exploited for therapeutic gain is an active area of research.

## Materials and Methods

### Cell Culture

The HN12, HN31, and HN8 cell lines were cultured in DMEM + 10% FBS. The JHU-022 cell line was cultured in DMEM/F12 + 10% FBS. These cell lines are from lymph node metastasis derived from the oral cavity (38,39). TerT-Immortalized Normal Oral Keratinocytes (NOKs) were cultured in keratinocyte medium (Sciencell). Cells were grown in a humidified atmosphere of 5% CO_2_ at 37 °C. All cell lines were authenticated via Short Tandem repeat profiling (ATCC). Tert-Immortalized NOKs were a gift from the lab of Michael McVoy. Cells were screened for mycoplasma using PCR or MycoStrip^Tm^ mycoplasma detection kit (Invivogen).

### Viral packaging infection and preparation of FPN expressing Cell lines

The tet-inducible lentivirus vectors pLVX-TetOne-puro vector and pLVX-Tetone-puro-Lucif control vector were purchased from Takara Bio. The FPN protein coding sequence was amplified via PCR to have sequences complimentary to the pLVX-TetOne vector on the 5’ and 3’ termini. The pLVX-TetOne-puro vector was digested with *BamHI* and *EcoRI* and combined with the FPN CDS using NEB HiFi DNA assembly master mix (New England Biolabs) to create the pLVX-tet-FPN vector. Clones were screened and confirmed via sequencing.

Viral particles were created via co-transfection of the pLVX-tet-FPN vector and 4^th^ generation packaging vectors using the Lenti-X packaging Single Shots (VSV-G) (Takara Bio) in Lenti-X HEK 293T cells. Viral particles for the luciferase control plasmid were created similarly. Two days after transfection, viral particles were collected from supernatant and filtered with a 0.45 μm filter. The HN12, JHU-022, and NOK cell lines were transduced with viral particles in the presence of polybrene (10 μg/mL) at ~50-70% confluency. After 48 hours, cells were passaged, and stable clonal populations were selected using puromycin.

### Cell growth

Cell growth and proliferation was measured by cell counting or CellTiter blue assay (Promega). For cell counting, cells were seeded in 6-well plates at 25,000 cells per well and allowed to grow for 4 days. After incubation, cells were harvested and counted using hemocytometer. For growth curves, cells were seeded in a clear-bottom, black 96 well plate at 1000 cells per well. Cell growth was assessed at 24, 48, 72, and 96 hours using 10 μL CellTiter-blue reagent and measuring fluorescence (560nm Ex/590nm Em) after 1 hour.

### Cell viability/survival assays

CellTiter-glo assays were performed by seeding cells at 3-5k cells per well (such that cells were at least 30% confluency at time of treatment) in a black 96 well plate. Supplemental compounds were added at the desired final concentration the next day and incubated for 3 days. After incubation, 25 μL of CellTiter-glo reagent was added to each well and measured on a H1 BioTek plate reader as per manufacturer’s protocol. Non-treated cells seeded in parallel were used to determine total cell growth.

Cell survival assessed via flow cytometry utilized 7-AAD staining. Cells were seeded in 6 well plates and compounds were added at the final concentration the next day and incubated for 3 days. Cells were trypsinized and combined with cell culture supernatant. Cells were pelleted and resuspended in PBS + 10% FBS and stained with 7-AAD (ThermoFisher). 7-AAD staining was assessed using a BD FACSCanto II cell cytometer with 488nm excitation and emission was collected using a 670LP filter. Data was analyzed using BD FACSDiva 8.0.1 software.

### Clonogenic assays

Cells were harvested and seeded in a 12 well plate at 250 cells per well. Doxycycline was added at a final concentration of 0.5 μg/mL. Plates were incubated for 7-10 days (until colonies >~50 cells began to form). Afterwards cells were washed and fixed using methanol. To each well, 0.1% crystal violet was added and wells were washed and colonies were counted.

To determine if FPN mediated inhibition of cell growth was transient or permanent, FPN-induced and control cells were grown for 3 days with doxycycline. Cells were then washed and doxycycline free media was added and cells were grown for an additional 2 days. After incubation, cells were trypsinized and seeded at 250 cells per well in a 12 well plate and incubated for 7-10 days. Colonies were then counted as described above.

### Western Blot

Cells were lysed in RIPA lysis buffer (150 mM NaCl, 1% TritonX-100, 0.5% Deoxycholic acid, 0.1 % SDS, 50 mM Tris pH 7.4) plus protease inhibitor cocktail (Millipore-Sigma). Samples were separated on 4-12% Bis-Tris PAGE gels and transferred to PVDF membranes set in NuPAGE transfer buffer (Thermo-Fisher) containing 20% methanol and blocked using 5% milk in TBST. Membranes were probed with primary and secondary antibody and detected using chemiluminescence in SynGene G-Box. Antibodies used: GAPDH (Novus Bio - #NB100- 56875), FPN (Novus Bio - #NBP1-21502), TFR1 (ThermoFisher - #13-6800), FTH1 (Cell Signaling Technology - #4393), NDGRG1 (Cell Signaling Technology - #9485), p21 (Cell Signaling Technology - #2947).

### Real Time qRT-PCR

The RNeasy kit (Qiagen) was used to purify RNA from cells. Residual DNA was removed using DNA-free DNAse kit (ThermoFisher). The cDNA was generated using the high capacity cDNA Reverse transcriptase kit (ThermoFisher). Real Time qRT-PCR was performed using SYBR-green based detection on a Quant Studio 3 real time PCR system using PowerUP Syber green MasterMix (ThermoFisher). Primers for GAPDH and beta-actin were used as endogenous controls to which RNA levels of target genes were normalized to. Primers used for qRT-PCR analysis are listed in supplementary table S1.

### Labile Iron pool assay

Labile iron levels were measured using the FerroOrange labile ferrous iron detecting probe (Goryo Chemical Inc). Cells were grown in a 6 well plate for 72 hours plus/minus doxycycline. Cells were then seeded in a black, clear bottom, 96 well plate at 5000 cells per well. The next day, cells were washed with serum free media containing 50 μM DFO and twice with HBSS to remove free extra-cellular iron. Cells were then incubated with 1μM of probe in HBSS for 30 minutes at 37 °C. After incubation, probe was removed from the cells and fresh HBSS was added, and fluorescence (Ex 542, Em 572) was measured. Values were normalized to the amount of total cells as measured by CellTiter-glo assay.

### Cell cycle analysis

Cells were grown with doxycycline for 3 days (HN12) or 4 days (JHU-022, NOKs) to induce FPN expression. Cells were harvested by trypsinization and washed with PBS and resuspended in 300 μL of PBS and fixed via drop wise addition of −20 °C absolute ethanol to a final concentration of 70% and incubated at −20 °C for at least 1 hour. Fixed cells were washed with PBS and resuspended with FxCycle PI/RNase staining solution (ThermoFisher #F10797). Cells were incubated for at least 30 minutes at room temperature. Fluorescence was measured on a BD FACSCanto II cell cytometer using 488nm excitation and emission was collected using a 670LP filter. FlowJo v10.8.1 was utilized to perform cell cycle calculations.

### Edu Incorporation

Edu incorporation was measured using the Click-it Plus Edu Alexa Fluor 647 Flow Cytometry Assay Kit (ThermoFisher C10634). Cells were grown for 3 days with doxycycline to induce trans gene expression and were then incubated with 20 μM Edu for 1.5 hours. After Edu incubations, cells were washed and trypsinized. Incorporated Edu was labeled as per manufacturer’s protocol. After labeling Edu, total nuclear DNA content was stained using FxCycle PI/RNase solution. Fluorescence was measured on a BD FACSCanto II cell cytometer using 633 excitation and emission was collected using a 660nm/20BP filter. PI fluorescence was measured as detailed above. FlowJo v10.8.1 was utilized to perform Edu incorporation calculations.

### Beta-galactosidase activity assay

The Beta-galactosidase activity assay was purchased from Abcam (Ab287846). Cells to be tested for activity were grown in a 6 well plate for 7 days in the presence of doxycycline. For input, 500,000 cells were used and protocol was followed per manufacturer’s instructions. Reaction was performed at room temperature in a 96 well plate and read using a H1 BioTek plate reader.

### Phospho-(S139) gamma-H2Ax immunofluorescence

Cells were seeded into a 4-chamber slide (Millipore-Sigma) and grown for 72 hours in 0.5 μg/mL doxycycline. Cells were fixed in 4% paraformaldehyde for 10 minutes and cell membrane was permeabilized using 90% methanol for 10 minutes. Slides were blocked using 1% BSA and incubated using anti-p-γH2Ax antibody (Abcam #242296) for 1 hour at room temp then stained using FITC conjugated secondary antibody. Coverslips were mounted using Prolong Anti-fade with DAPI (ThermoFisher) and imaged on a Keyence BZ-X810 microscope.

For calculations, staining was repeated three times. For each biological replicate, 100-150 random cells were imaged and counted and cells positive for p-γH2Ax staining were quantified against the total amount of cells.

### Mouse Xenograft Studies

All animal procedures were approved by the VCU Institutional Animal Care and Use Committee (IACUC) under protocol AD10002926. Six week old female NSG (NOD-*scid* IL2Rgamma^null^) mice were purchased from the VCU Massey Cancer Mouse Models Core. To each mouse, 50,000 HN12-FPN cells in serum free media were injected into the left cheek in a 1:1 mixture of cells to Matrigel. Five days after injection mice were randomized into two groups. FPN expression was induced in mouse xenografts by adding 2mg/mL of doxycycline to the drinking water of one group. The control group received normal drinking water. Seven mice for each group (plus/minus doxycycline) were used for a total of n=14 mice. After 21 days, mice were sacrificed, and tumor volume was recorded, and tumors were weighed. No surviving mice were excluded from the experiment. Differences in groups were calculated using a one-tailed T-test of equal variances.

### Statistical analysis

All *in vitro* experiments were performed at least 3 times. Statistical analysis was performed in excel. Comparison tests were performed using a two-tailed unpaired T test of equal variance. Results are reported as mean plus/minus the standard deviation of the mean.

## Acknowledgements

Funding for this work was provided by NIH grants F32DE029412 (BRB) and R01DE023304 (JPL). We would like to thank the lab of Michael McVoy for providing the NOK cell line. We would like to thank Drs. Anthony Faber, Kostas Floros, Jinyang Cai, and Molly Bristol of the VCU Philips Institute for reagents, cell lines, and critique of the study.

Services and products in support of the research project were generated by the VCU Cancer Mouse Models Core Laboratory and the VCU Flow Cytometry Shared Resource, supported, in part, with funding from NIH-NCI Cancer Center Support Grant P30 CA016059. We would like to acknowledge Dr. Jennifer Koblinski for assistance developing the animal model and Dr. Bin Hu of the VCU Cancer Mouse Models Core for his technical help in this study.

## Author Contributions

**B. Ross Belvin:** Conceptualization, Investigation, Data Curation, Formal Analysis, Funding Acquisition, Methodology, Visualization, Writing – Original Draft, Writing – Review and Editing. **Janina P. Lewis:** Conceptualization, Supervision, Project Administration, Funding Acquisition, Methodology, Writing – Review and Editing.

## Supplemental Figure Legends

**Figure S1.**
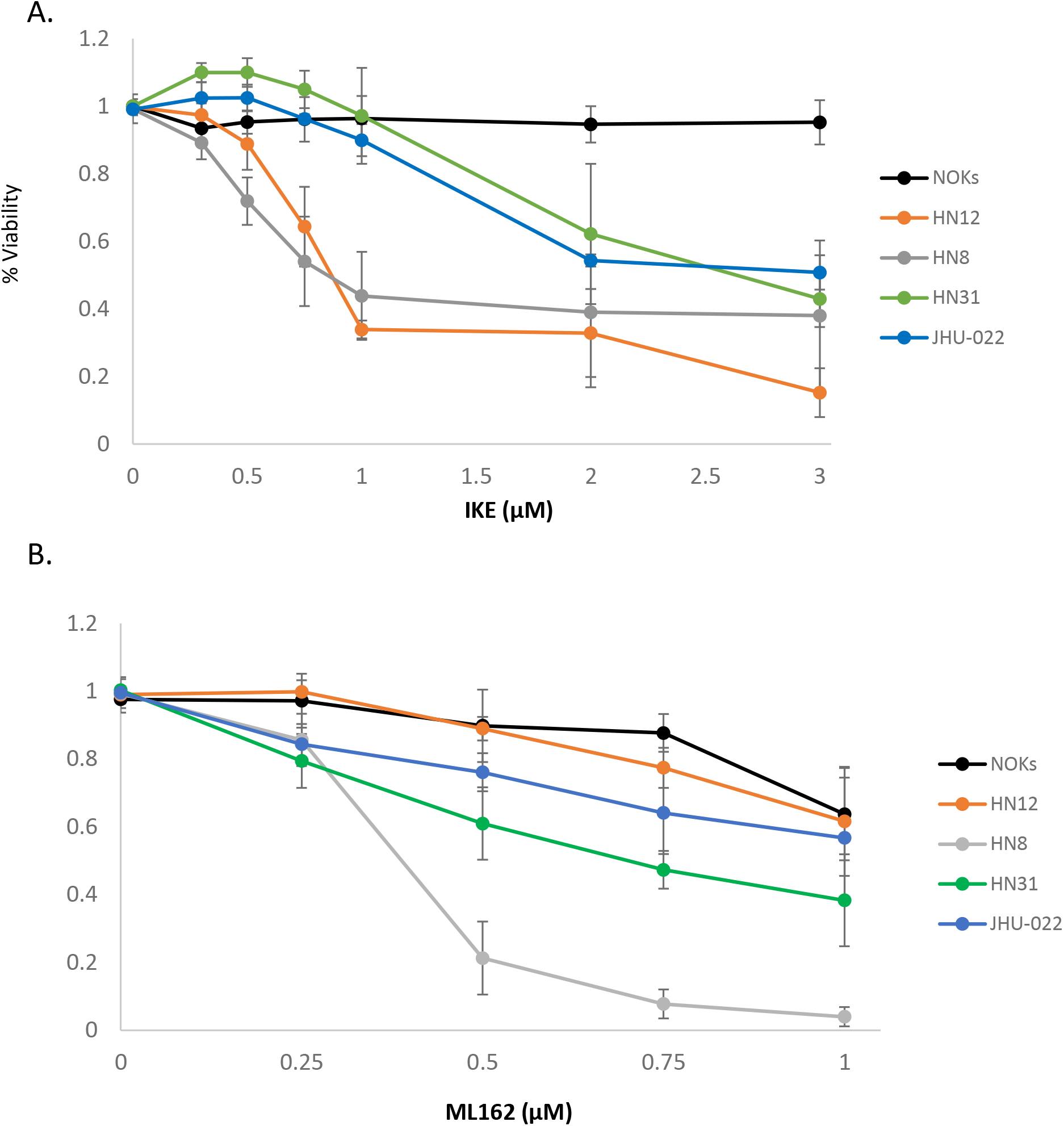
HNSCC cell lines are sensitive to ferroptosis inducers. Cells were treated with increasing concentrations (**A**.) Imidazole Ketone Erastin (IKE) or (**B**.) ML162 and cell survival was assessed by CellTiter-glo. Cells were treated for 72 hours with the indicated concentrations of drugs.

**Figure S2.**
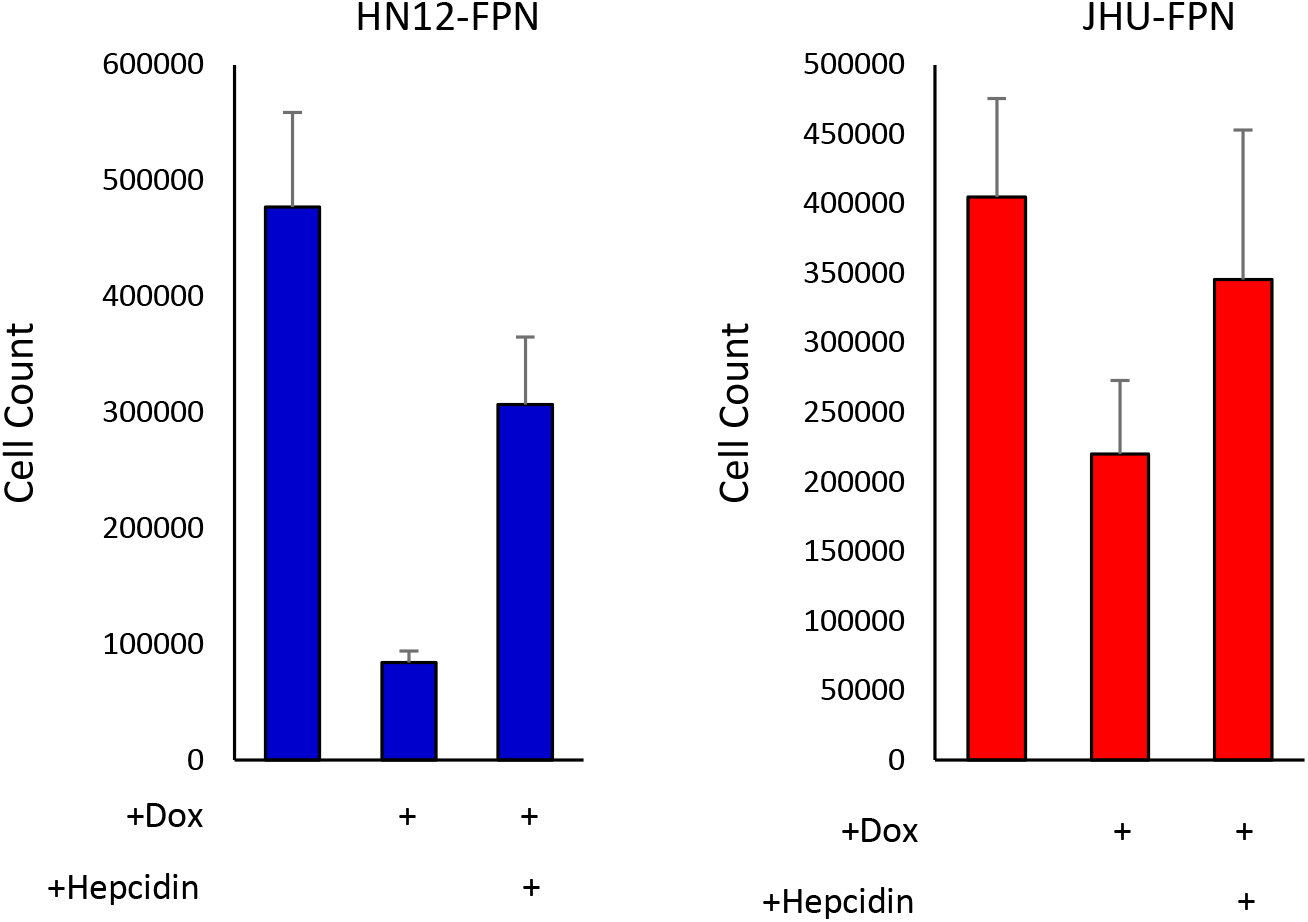
Hepcidin rescues growth of FPN expressing HNSCC cell lines. HN12-FPN (**A**.) and JHU-FPN (**B**.) were seeded in 6 well plates and treated with doxycycline (0.25 μg/mL), hepcidin (10 μM), or left untreated. Cells were grown for 4 days, after which cells were trypsinized and counted via hemocytometer.

**Figure S3.**
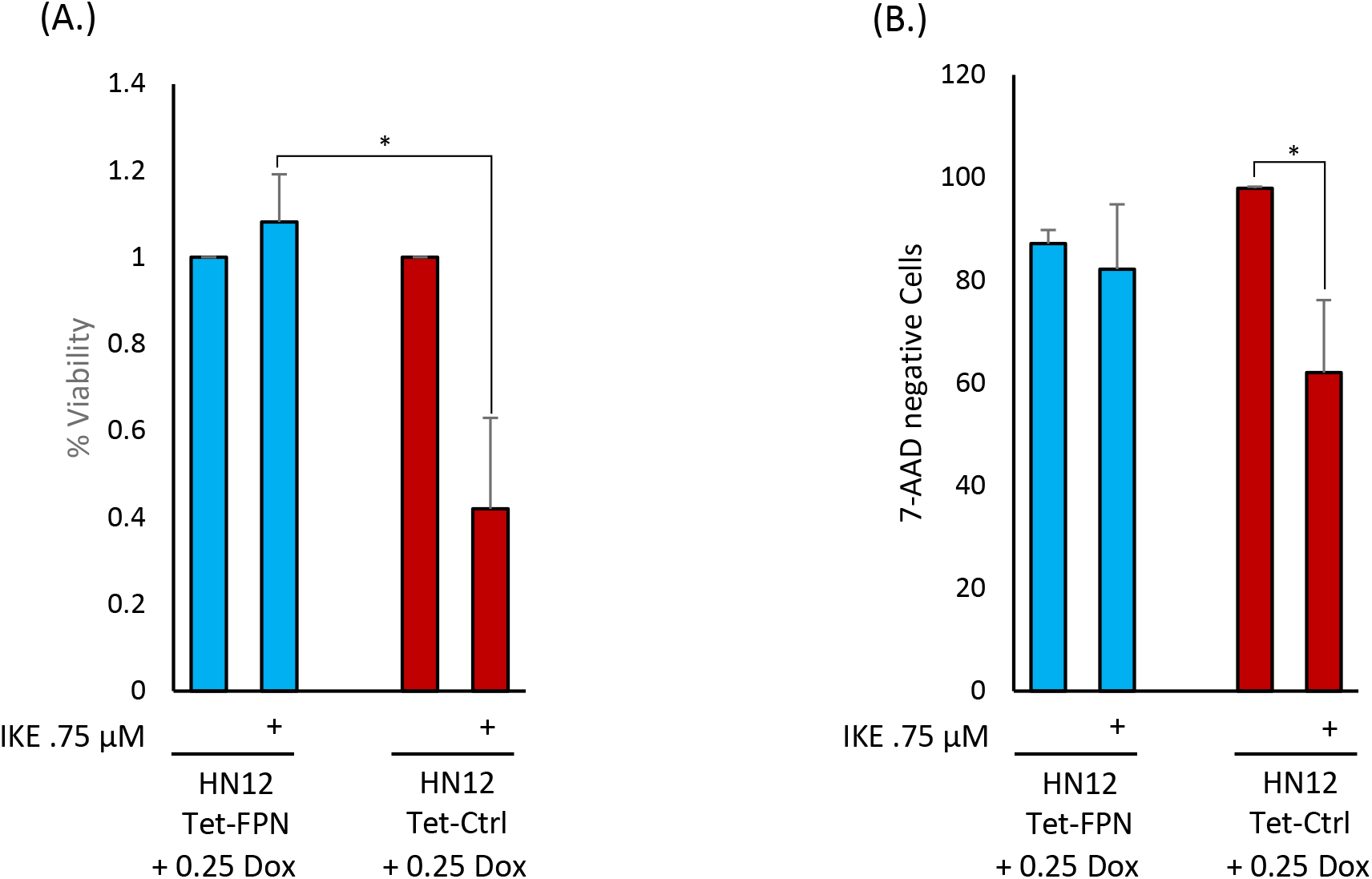
**A**. HN12-FPN cells or HN12-Luc cells were seeded in a 96 well plate +/-doxycycline (0.25 μg/mL). The next day IKE (0.75 μM) was added and cells were treated for 3 days. Cell viability was assessed via CellTiter-glo assay. IKE treated conditions were normalized to untreated conditions plus doxycycline. **B**. HN12-FPN and HN12-Luc cells were seeded 6 well plates plus 0.25 μg/mL doxycycline. The next day 0.75 μM IKE was added ad cells were incubated for 3 days. After incubation cells were trypsinized and stained with 7-AAD to assess cell survival. Results were analyzed via flow cytometry.

**Figure S4.**
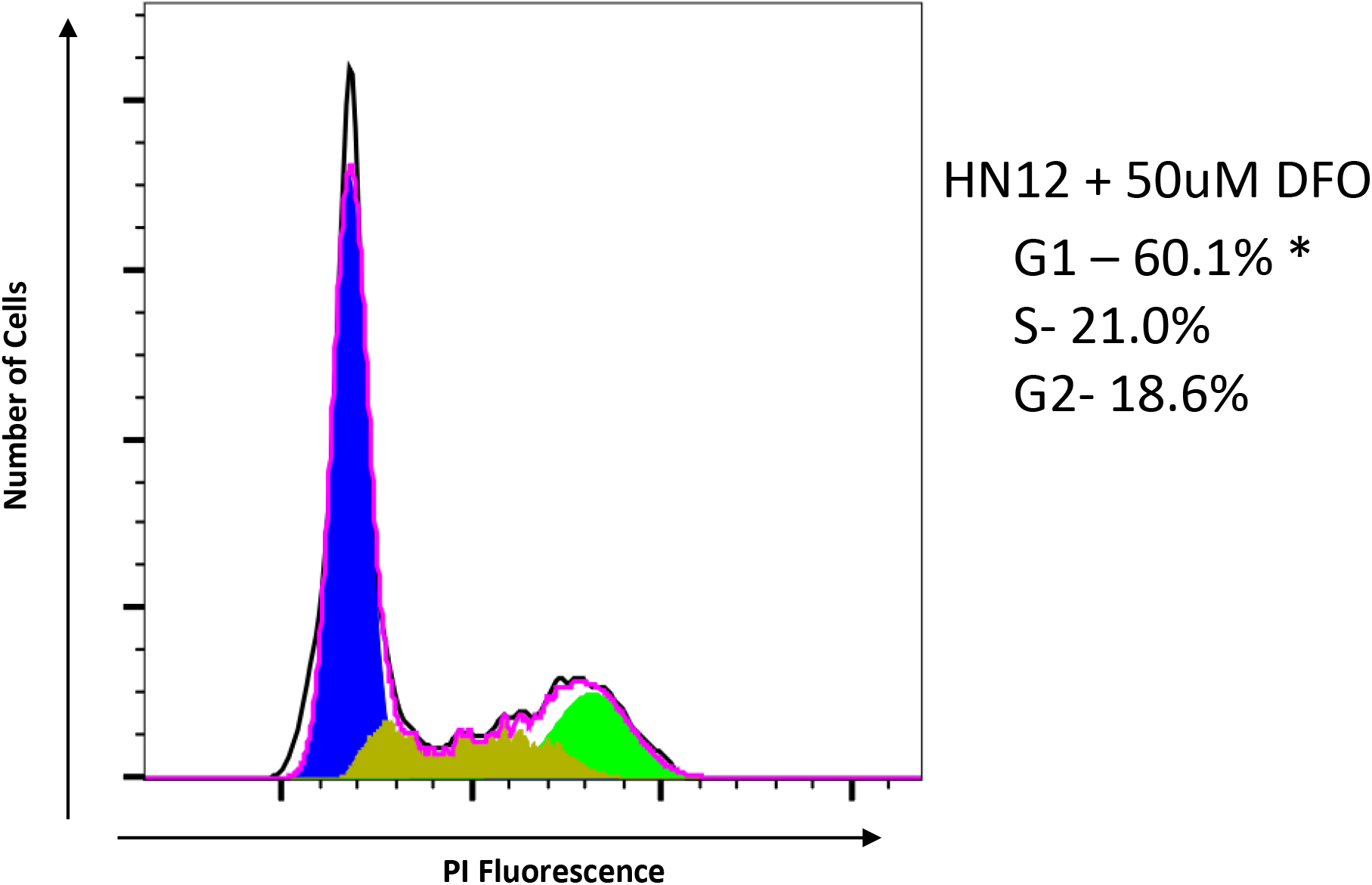
DFO arrests cells in G_0_/G_1_ phase. HN12 cells were treated with 50 μM for 72 hours. Cells then stained with propidium iodide to measure DNA content and measured using flow cytometry.

**Figure S5.**
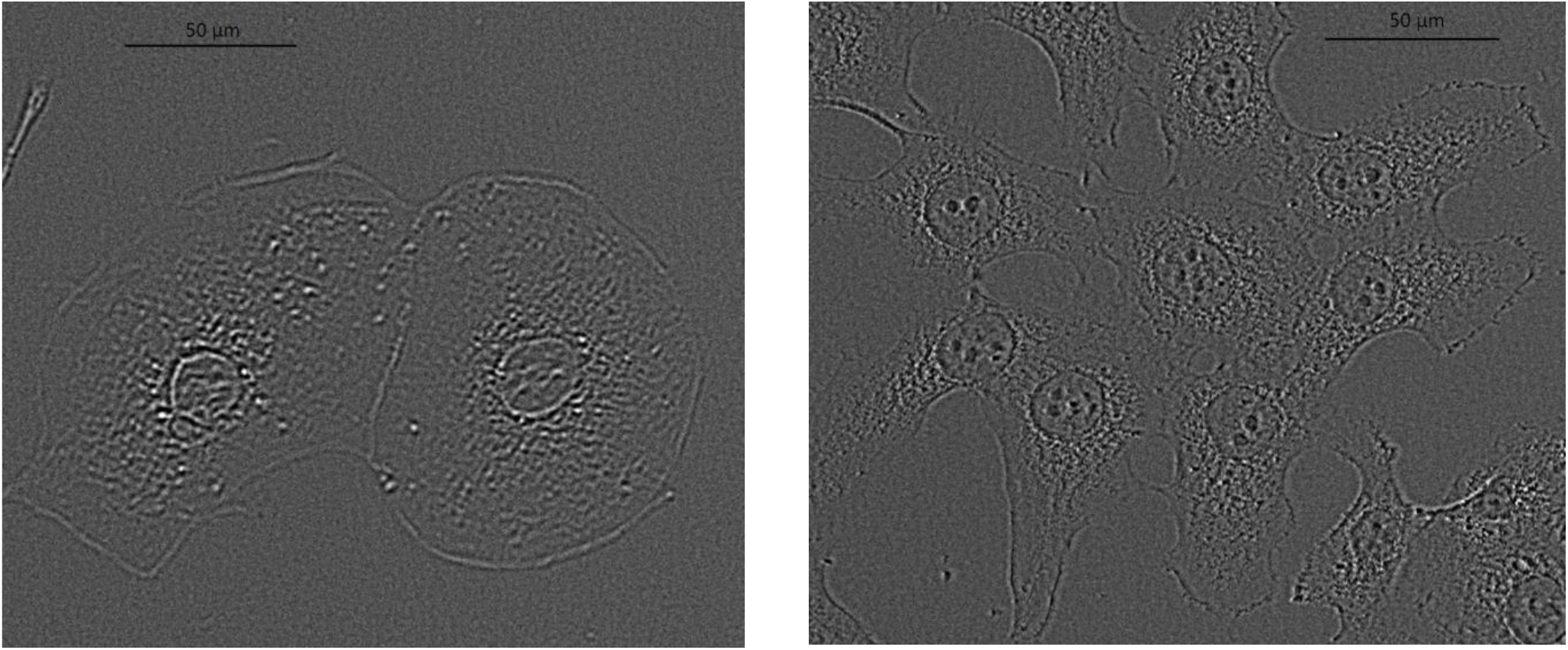
HN12-FPN cells have typical senescent cell morphology. Phase contrast images of HN12-FPN (**A**.) and HN12-Luc (**B**.)cells stained for γH2Ax.

**Figure S6.**
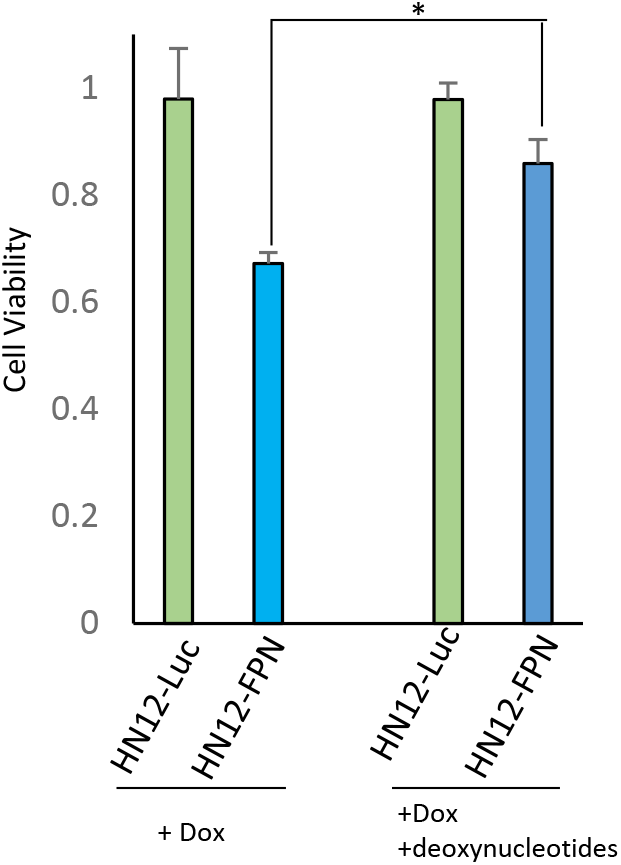
The addition of deoxynucleotides partially rescues cells from growth inhibition. HN12-FPN and HN12-Luc cells were seeded in 96 well plates and grown with 0.25 μg/mL of doxycycline. Cultures were supplemented with 10 μg/mL of 2’-deoxyadenosine, 2’-deoxycytidine, 2’-deoxyguanidine, and thymidine where indicated. Cells were cultured for 48 hours and growth was assessed using Celltiter-blue assay. Growth was normalized to cells untreated with doxycycline.

