## Supplementary figures and images for "Ferroportin Depletes Iron Needed for Cell Cycle Progression in Head and Neck Squamous Cell Carcinoma"

### Supplemental Figures

#### Supplementals

##### Figure S1

A.

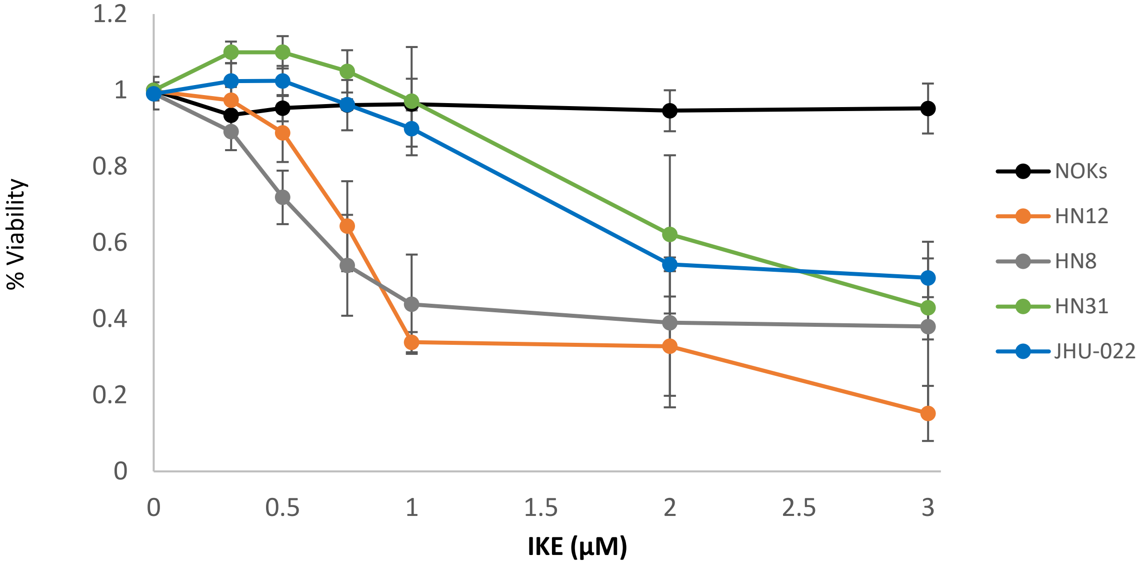


B.

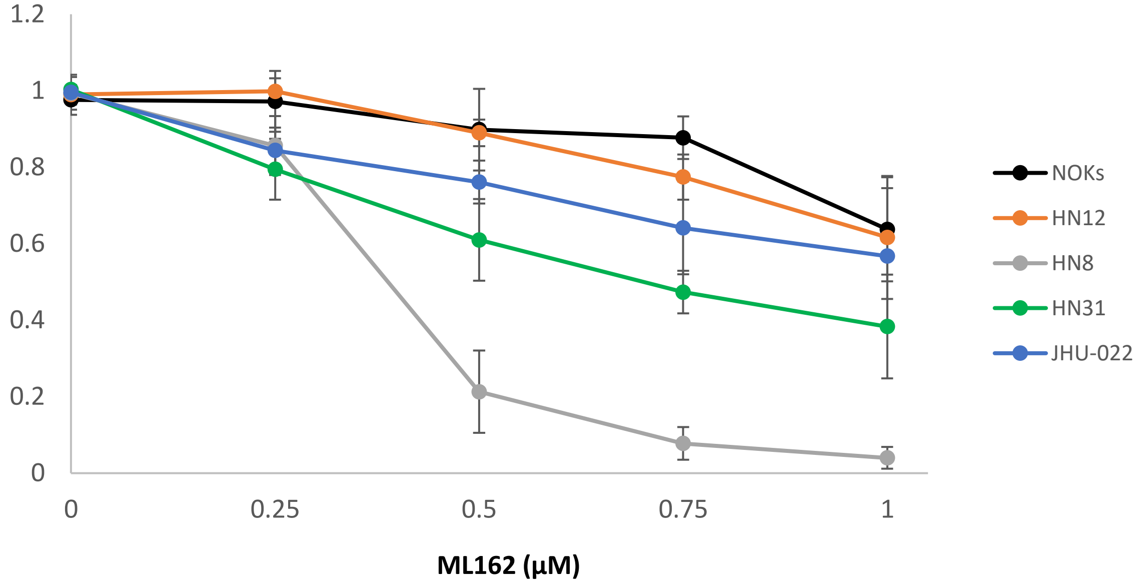

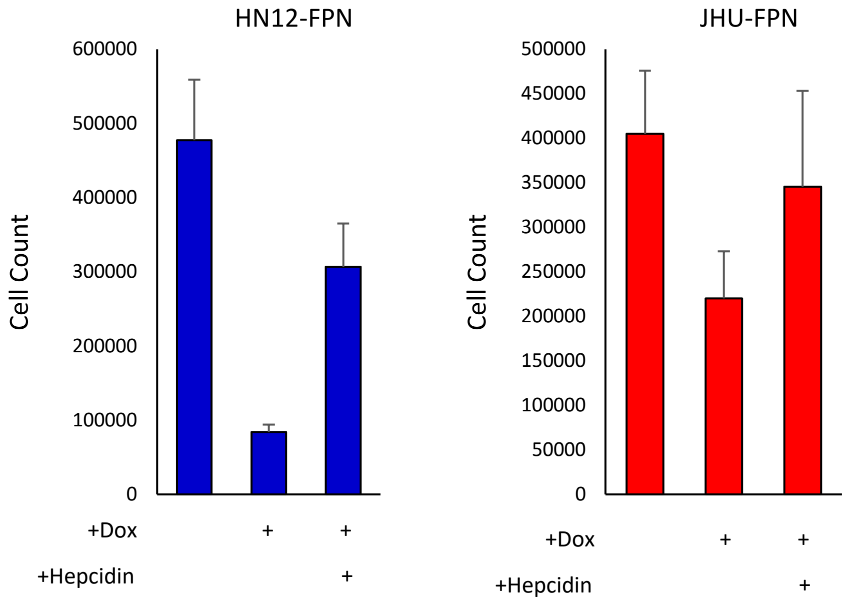

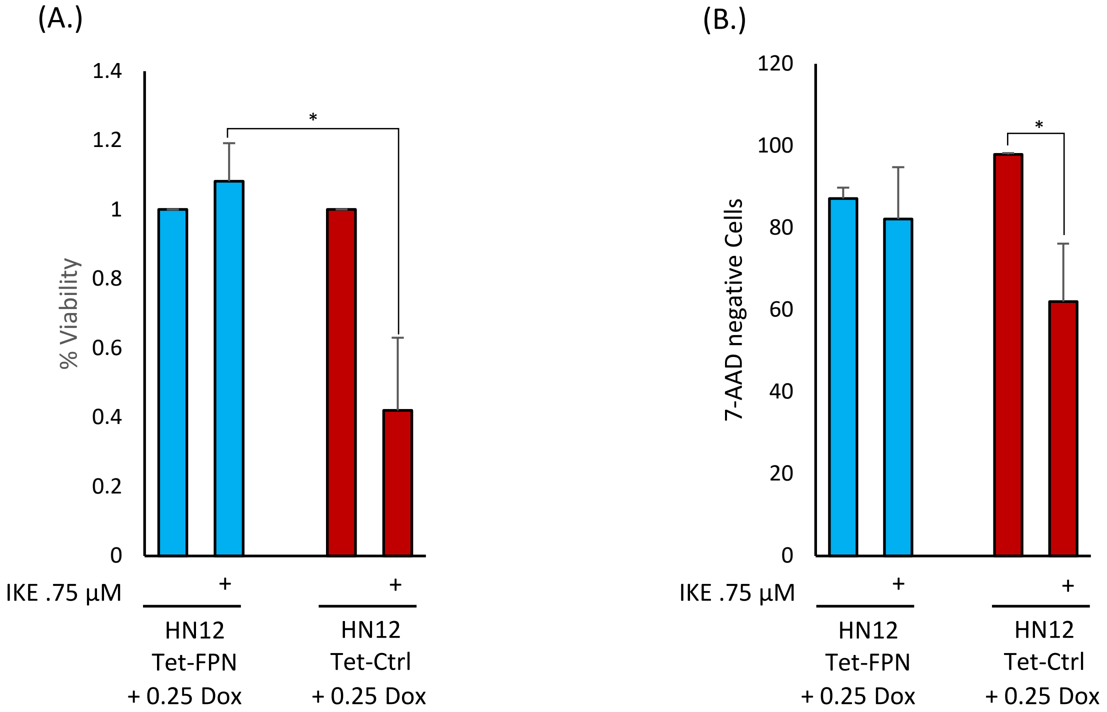

Figure S4

`

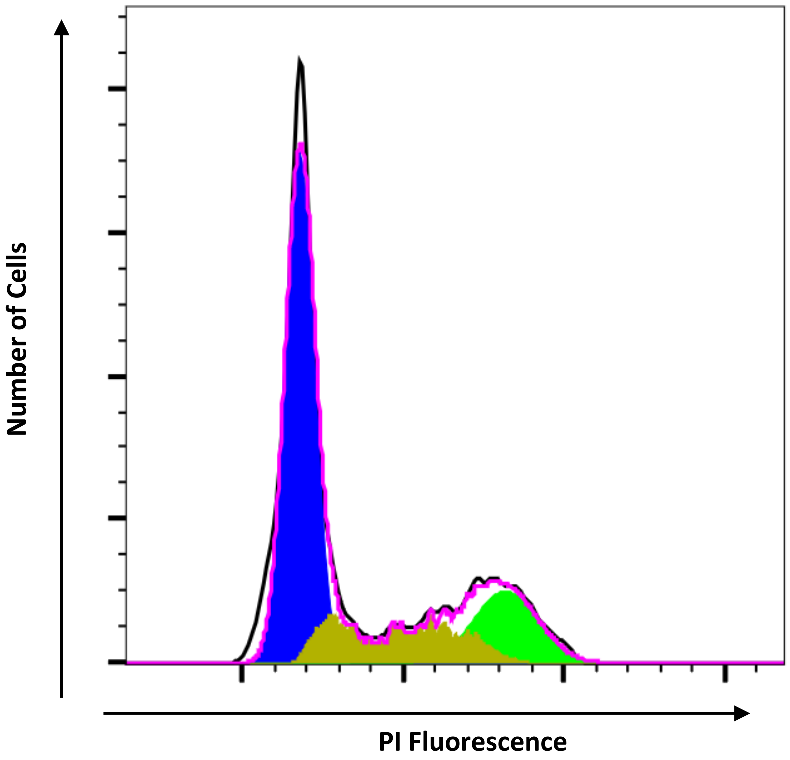


HN12 + 50uM DFO

G1 – 60.1% *

S- 21.0%

G2- 18.6%

Figure S5

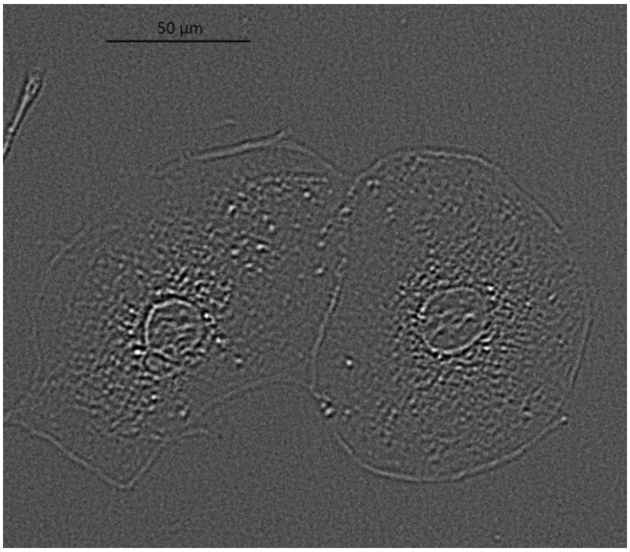


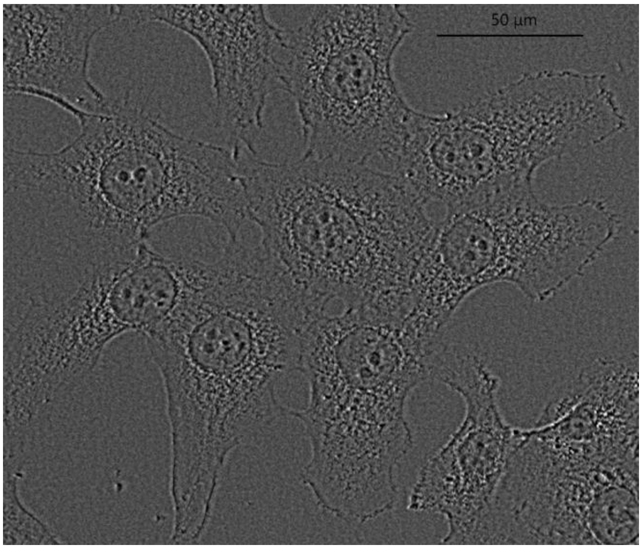

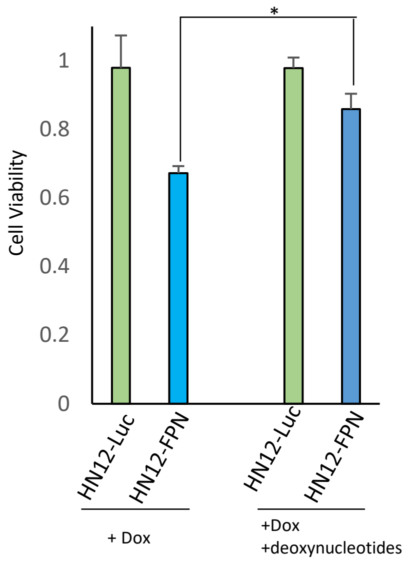
