## Supplemental Table 1 for "Ferroportin Depletes Iron Needed for Cell Cycle Progression in Head and Neck Squamous Cell Carcinoma"

| Primer Name | Primer Sequence |
|---|---|
| p21-qPCR-F | CGC TCT ACA TCT TCT GCC TTA GTC |
| p21-qPCR-R | GAA CCT CTC ATT CAA CCG CCT AG |
| CyclinA-qPCR-F | TGG TTA GTT GAA GTA GGA GAA G |
| CyclinA-qPCR-R | TTG GTG TAG GTA TCA TCT GTA ATG |
| CyclinB-qPCR-F | TGG TTG ATA CTG CCT CTC C |
| CyclinB-qPCR-R | TCT GAC TGC TTG CTC TTC C |
| CyclinD-qPCR-F | TGA ACT ACC TGG ACC GCT TC |
| CyclinD-qPCR-R | AGC TTG TTC ACC AGG AGC AG |
| B-actin-qPCR-F | TTG CCG ACA GGA TGC AGA AGG A |
| B-actin-qPCR-R | AGG TGG ACA GCG AGG CCA GGA |
| GAPDH-qPCR-F | TGG TAT CGT GGA AGG ACT CAT GAC |
| GAPDH-qPCR-R | ATG CCA GTG AGC TTC CCG TTC AGC |
| FPN-pLVX-Tetone-F | CCC TCG TAA AGA ATT ATG ACC AGG GCG GGA GAT CAC AAC |
| FPN-pLVX-Tetone-R | GAG GTG GTC TGG ATC TCA AAC AAC AGA TGT ATT TGC TTG |
